# Genome regulation and gene interaction networks inferred from muscle transcriptome underlying feed efficiency in Pigs

**DOI:** 10.1101/2020.03.20.998203

**Authors:** Victor AO. Carmelo, Haja N. Kadarmideen

## Abstract

Improvement of feed efficiency (FE) is key for sustainability and cost reduction in pig production. Our aim was to characterize the muscle transcriptomic profiles in Danbred Duroc (Duroc) and Danbred Landrace (Landrace), in relation to FE for identifying potential biomarkers. RNA-seq data was analyzed employing differential gene expression methods, gene-gene interaction and network analysis, including pathway and functional analysis. We compared the results with genome regulation in human exercise data. In the differential expression analysis, 13 genes were differentially expressed, including: *MRPS11, MTRF1*, TRIM63, MGAT4A, KLH30. Based on a novel gene selection method, the divergent count, we performed pathway enrichment analysis. We found 5 significantly enriched pathways related to feed conversion ratio (FCR). These pathways were mainly mitochondrial, and summarized in the mitochondrial translation elongation (MTR) pathway. In the gene interaction analysis, highlights include the mitochondrial genes: PPIF, MRPL35, NDUFS4and the fat metabolism and obesity genes: *AACS, SMPDL3B, CTNNBL1, NDUFS4* and *LIMD2*. In the network analysis, we identified two modules significantly correlated with FCR. Pathway enrichment of modules identified MTR, electron transport chain and DNA repair as enriched pathways. In the network analysis, the mitochondrial gene group *NDUF* was a key hub group, showing potential as biomarkers. Comparing with human transcriptomic exercise studies, genes related to exercise displayed enrichment in our FCR related genes. We conclude that mitochondrial activity is a driver for FCR in muscle tissue, and mitochondrial genes could be potential biomarkers for FCR in pigs. We hypothesize that increased FE mimics processes triggered in exercised muscle.

## Introduction

In commercial pig production, the cost of feed is the highest individual economic factor (Jing, Hou et al. 2015, Gilbert, Billon et al. 2017). Furthermore, reduction in feed consumption per unit growth is beneficial for the environment, which is a key factor in being able to maintain sustainable and resource efficient production. In this context, there have been continuous efforts to increase feed utilization efficiency in pigs through selective breeding. In the Danish Production pig population, breeding is done at a core central facility where potential breeding sires are tested for FCR through accurate individual measurements of feed intake and growth. Danish production pigs are crossbreds, with the maternal line being Landrace x Danbred Yorkshire, and the paternal line being Durocs The Durocs are well-known for being heavily selected for growth and efficiency, while the two other breeds have had more heavy selection on litter size or piglet survival related traits.

Feed efficiency can be defined in several ways, with the main ones being Residual Feed Intake RFI(Koch 1963) and FCR. FCR is the ratio between feed consumed and growth, while RFI is based on the residual between predicted feed intake and actual feed intake given growth. In general, it is reported that selection for low FCR will result in co-selection for related traits, namely growth rate and body composition (Nkrumah, Basarab et al. 2007, Gilbert, Billon et al. 2017, Yi, Li et al. 2018). In contrast, selection for RFI is more directly focused on metabolic efficiency irrespective of daily gain and growth (Nkrumah, Basarab et al. 2007, Gilbert, Billon et al. 2017, Yi, Li et al. 2018). In general, RFI and FCR are strongly correlated, with a correlation above 0.7 and both show low to medium heritability(Do, Strathe et al. 2013). In general, FCR is simpler to calculate, as RFI calculation is dependent on individual population and production factors (Hoque, Kadowaki et al. 2009, Do, Strathe et al. 2013). However, in pig production, the side effects of FCR selection and simplicity are desired traits, thus perhaps explaining why the pig population in Denmark and in general pig production, FCR has been the main efficiency phenotype used for selection (Gilbert, Billon et al. 2017). One can also hypothesize that FCR is more easily translatable between breeds/populations, as it is a simple dimensionless ratio, which has a simple and generally comparable interpretation. In contrast, it is more difficult to easily compare RFI values across different populations or breeds. In regards to the biological and/or genetic background of FCR in pigs, the results remain somewhat elusive(Ding, Yang et al. 2018), thus inviting for further analysis on the topic.

The key tissue in pig production is muscle, as pig carcasses are valued according to lean meat content. Skeletal muscle is a key organ in carbohydrate and lipid metabolism and plays a large part in the storage of energy from feed (Turner, Cooney et al. 2014, Morales, Bucarey et al. 2017), especially as lean growth has been one of the main goals of pig breeding programs. Increased efficiency has also been positively associated with various meat quality parameters (Czernichow, Thomas et al. 2010, Lefaucheur, Lebret et al. 2011, Smith, Gabler et al. 2011, Faure, Lefaucheur et al. 2013, Horodyska, Oster et al. 2018), showing that improved FE can have multiple positive outcomes. There are only a few studies analyzing muscle tissue transcriptome pf pigs in a FE context(Jing, Hou et al. 2015, Vincent, Louveau et al. 2015, Gondret, Vincent et al. 2017, Horodyska, Wimmers et al. 2018), and thus our knowledge of the muscle transcriptomic background of FE is somewhat limited. In general, the studies available have relied on small samples sizes, weak statistical thresholds and categorical division of lines divergently selected for FE. This means that more studies are still needed to uncover the true underlying transcriptomic background of FE in muscle tissue.

Here, in our study, we aim to characterize the transcriptomic profiles and link them to FE traits measured in Duroc and Landrace, purebred pigs, by fitting FE as a continuous trait over a full spectrum of efficiency, from high to low. Furthermore, the pigs selected for the study all came out of the potential breeding sire population, with no pigs negatively selected for FE, thus better representing real world breeding scenarios than using negative FE selection. We analyzed the muscle transcriptome based on several layers of statistical-bioinformatics analyses: differential expression (DE), gene-gene interaction and network analysis, which was followed up by pathway and functional analysis. The rationale behind the approach was to reveal potential biomarkers that are functionally important and are predictive of FE in pigs. Dealing with complex yet subtle phenotypes can be a challenging, as the signal to noise ratio can be high, and it may be impractical or costly to collect large sample sizes. Therefore, we also suggest a novel method for selecting features based on overall p-value distributions, the divergent count.

To gain more insight on the molecular and functional background of FE, we also hypothesized, that the mechanism between differences in the muscle transcriptome of breeds with different efficiency could be similar to the differences between a rested and an exercised muscle, We adapted a translational genomics approach to investigate this, comparing human data with our data.

## Materials and Methods

### Sampling and Sequencing

In total, 41 purebred male uncastrated pigs where sampled for this study from two breeds, with 13 Danbred Durocand 28 Danbred Landrace pigs. All pigs were raised at a commercial breeding station at Bøgildgard owned by the pig research Centre of the Danish Agriculture and Food Council (SEGES). The pigs where raised from ∼7kg until ∼100kg at the breeding station. During this time, all feed intake was measured starting at 28kg and for a period of 40-70 days based on the viability of each pig. All pigs were routinely weighed several times, including at testing start and end for calculation of FCR. FCR was calculated by dividing the growth in the testing period with the feed consumption. Residual Feed Intake (RFI) was also estimated based on the residuals of the following model, from Do et al(Do, Strathe et al. 2013):

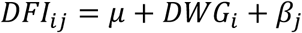

Where DFI is daily feed intake and DWG is daily weight gain in the period, and β is the batch effect. RFI was calculated separately for each breed, and based on data from a larger population (Duroc *n*=59 and Landrace n=50).

Muscle tissue samples from the psoas major muscle were extracted immediately post slaughter and preserved in RNAlater (Ambion, Austin, Texas). Sample were kept at −25 C, as per protocol, until sequencing

#### Sequencing

Sequencing was done on BGISEQ-500 platform using the PE100 (pair end, 100bp length) with Oligo dT library prep at BGI Genomics.

### QC, Mapping and Read Quantification

Reads were trimmed and adapters removed using Trimmomatic (Bolger, Lohse et al. 2014) version 0.39 with default setting for paired end reads. The QC on the data was done both pre- and post-trimming using FastQC v0.11.9(. The reads were mapped using STAR aligner(Dobin, Davis et al. 2013) version 2.7.1a using default parameters with a genome index based on sus scrofa version 11.1 and using ensemble annotation *sus scrofa* 11.1 version 96 for splice site reference. Default parameters were used for mapping except for the addition of read quantification during mapping using the --quantMode GeneCounts setting. All statistic for the reads can be found in supplementary data 1.

### Differential Expression Analysis

To analyze the relationship between FCR and gene expression, we applied the following overall model, and implemented it using several different methods:

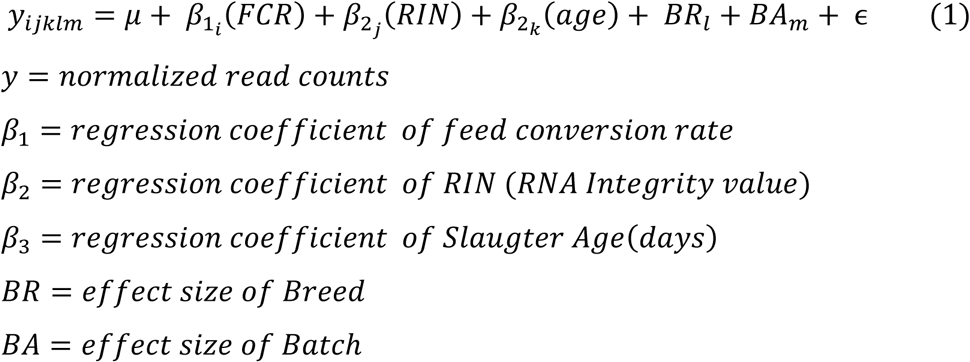

RNA integrity value (RIN) should be corrected for, as it affects expression, and the most appropriate way to correct this is to include it in the model(Gallego Romero, Pai et al. 2014). As the samples had different slaughter days, which affected the collection conditions, we also deemed it necessary to correct for this via the batch effect. Finally, we correct for Breed and age at slaughter, as these are biological factors, which can cause differences in expression.

We used the following 3 methods for the DEA: Limma (Ritchie, Phipson et al. 2015), edgeR (Robinson, McCarthy et al. 2010) and Deseq2(Love, Huber et al. 2014). This was done to increase the robustness of our analysis, as our phenotype of interest is expected to have a subtle effect on the transcriptome due to the complex nature of FE. In addition, we also fit the model for each breed separately using Deseq2, just removing the Breed as a covariate.

### Deseq2

We used Deseq2 version 1.22.2. In the Deseq2 analysis, the counts were filtered a priori requiring a minimum of 5 reads for each sample, resulting in a total of 10765 out of 25880 genes being included in the DE analysis in the common breed analysis, and 10687 and 11107 in Landrace and Duroc respectively. As the overall read counts were very similar across experiments (see supplementary data 1), it was deemed sufficient to filter pre normalizing. We then used the default analysis method based on our specified model.

### Limma

We used Limma version 3.38.3. For the Limma analysis, the counts were filtered based on the edgeR *filterByExpr*function and normalized using *calcNormFactors* from the same package, as suggested in the limma manual. This resulted in the inclusion of 11146 genes in the analysis. To fit the model we used the *eBayes* method in conjunction with our specified model.

### EdgeR

We used edgeR 3.24.3. We used the same normalization and filtering as in the Limma analysis, thus including the same number of genes. We used the *glmQLfit* function and *glmQLTest* to implement our model.

While we used to different set sizes in the analysis, this does not affect the results significantly, as the genes omitted in the Deseq2 analysis are all lowly expressed. Furthermore, in our further analysis we elected to use the smaller and more conservative Deseq2 set to become our reference set for selections and analysis. **Gene Pathway Analysis**

### Gene selection

To select a robust set of genes for a gene enrichment analysis when we have non-conservative p-value but only a limited number of genes with a FDR below 0.05, we applied the following strategy:

- Identify the overrepresentation of (low) p-values in comparison to a uniform p-value distribution in our data. We will call this the divergent count.
- Select the top N genes by p-value, where N is the estimated divergent count
- Among the top N genes, select those that are found in all three methods.

To find the divergent count D, we find the interval with the maximum positive divergence between our observed empirical p-values and the same number of uniformly distributed p-values. It is calculated as follows:

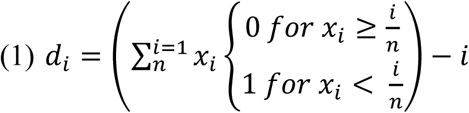

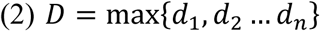

Where n is the total number of p-values, *x*_*i*_ is the i’th observed p-value in increasing order. Here i is both the index for x and the expected number of p-values between 0 and 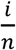 given a uniform distribution. D is the final divergent count, which is the maximum over all possible values of *d*..

#### GOrilla

To perform gene enrichment in GOrilla (Eden, Lipson et al. 2007, Eden, Navon et al. 2009), we translated our *sus scrofra* ensemble gene IDs into human ensemble gene IDs. The background set of genes used in GOrilla was the set of genes from the Deseq2 analysis. We used default settings. Furthermore, we used the Revigo (Supek, Bosnjak et al. 2011) analysis through GOrilla to generate summaries of our enrichment analysis, using default settings.

### Feed Efficiency measure

In this study, we elected to use weight gain/feed intake as our FCR measure. It fit the data better than RFI, and FCR is the metric used in the breeding program of our pigs.

### Pairwise Gene interaction Analysis

To continue our analysis of the top set of genes identified using the divergent counts in our DE analysis, we decided to apply a pairwise interaction model. First, we adjust the expression based on any factors and covariates that may affect expression for each gene. These factors are the same as in the general DE analysis, giving rise to the following linear model:

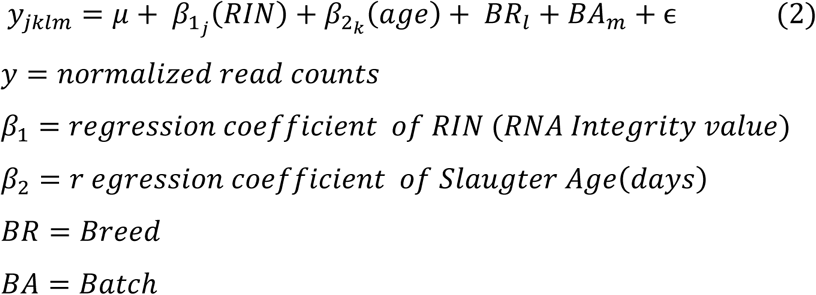

We then centered and scaled the residuals and then run a model for all pairwise gene interaction in our gene set. The reason we scaled and centered is that this leads to a more flexible and interpretable model regardless of the type of interaction. The interaction model was as follows:

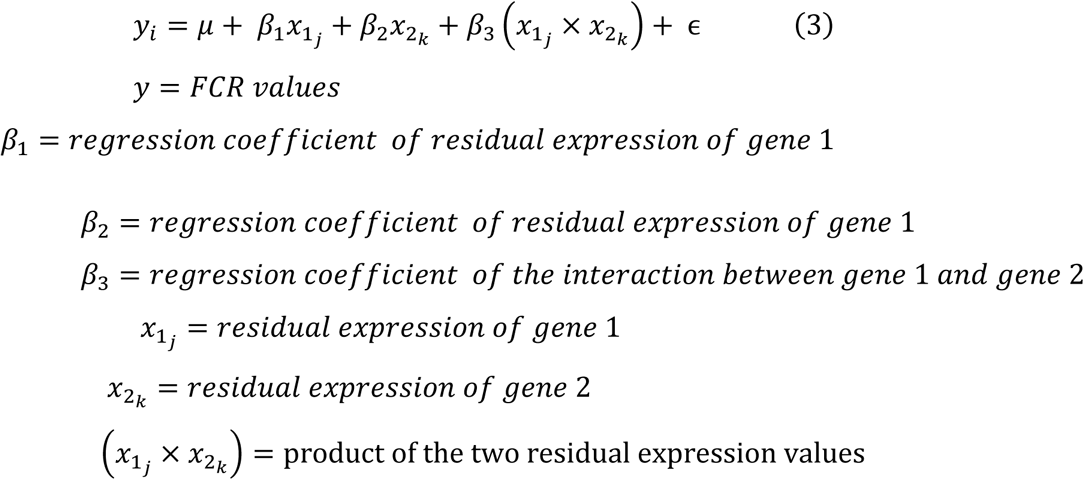

The next step was then to identify significant interactions. As the number of interaction in a dataset grows exponentially to the square of the input space, it is often difficult to detect effects based on classical multiple testing correction methods such as Bonferroni or FDR. This is especially true when dealing with complex phenotypes, as we generally do not expect to find individual large effects. Due to this, instead of focusing on individual results, for each gene, we calculated the divergent count, to assess the divergence of each genes distribution of interaction p-values. We then bootstrapped with replacement samples of 853 p-values from our empirical p-values 10^5^ times, calculating the divergent count each time, giving us a bootstrapped distribution of divergent counts, to compare with our empirical distribution

### Network analysis

To perform network analysis we used WGCNA(Langfelder and Horvath 2008). First, we filtered the read counts to only include genes with a minimum of 5 un-normalized reads, as was done for the Deseq2 analysis. We then created a correlation matrix based on all pairwise correlation in the data. The correlation matrix was based on un-normalized values as the correlation metric is based of comparison of the samples with themselves, thus it is not affected by the covariates. We then fit the ß parameter for the scaling of the network to create a scale free topology(Zhang and Horvath 2005). The scaled correlation matrix was used as an adjacency matrix that was used to generate the Topological Overlap Measures (TOM), which represents the final calculation of the relation between genes.

The TOM values of the genes where clustered using the *dynamicTreeCut* function from the dynamicTreeCut cut package with default setting, resulting in a number of module which are arbitrarily named based on colors.

The eigenvalue of each module was then calculated based on the normalized read counts and RIN adjusted count. We did these corrections in this step to remove the technical effects of library size differences and RIN from the eigenvalues, as we did not want technical effects to affect the eigenvalues.. The counts were normalized based on the *calcNormFactors* function from the edgeR package. After this, the counts were adjusted for RIN by fitting the following linear model: *expression* = μ + RIN + ϵ for all genes, and extracting the residual expression values. Highly correlating models where merged using the *mergeCloseModules* function using a default cut-off. We then calculated the Pearson correlation between corrected and normalized module eigenvalues and our traits and covariates. Pathway analysis was performed on the genes of highly correlated modules, with GOrilla and ReviGO as seen above. Finally, we also identified the top hub genes in relevant modules. This was done based on calculating the intramodular connectivity using the *intramodularConnectivity* function with default settings. We then selected the top hub genes base on the kWithin measure, which represents the connectivity within modules.

### Comparison to human exercise data

To test the hypothesis that differences in the muscle tissue transcriptome of Duroc and Landrace and/or FCR related genes mimic differences in rested and exercised muscle tissue, we compared our results with three human data sets(Murton, Billeter et al. 2014, Devarshi, Jones et al. 2018, Popov, Makhnovskii et al. 2019). For each data set, we performed the following:

1. Select the genes differentially expressed between breeds, based on the edgeR analysis
2. For FCR, use the 853 genes from divergent count set
3. Find the same set of genes in the human data – the breed/FCR matching genes. Genes are matched using the biomart R package, based on retrieving the external_gene_name of our sus scrofa ensemble gene identifiers.
4. Separate the human data into two parts – the breed matching set and the background set
5. Using a Fisher Exact test, compare the number of differentially expressed genes for the exercised vs rested muscle in the background set vs the breed matching set.
6. The steps for the breed were also applied to our divergent count set for FCR.

The reason edgeR was used in this part of the analysis, was because it was more flexible to fit to the publicly available data, allowing to compare our results to the other studies. As each dataset was formatted and analyzed differently, we had to process them individually. In the data set from Devarshi et al(dataset 1)(Devarshi, Jones et al. 2018), we chose to use the lean pre exercise vs lean post exercise group as our comparison, and significance was based on the reported cuffdiff analysis. For the set of Murton et al(dataset 2)(Murton, Billeter et al. 2014), we pooled all control vs exercise samples and analyzed them using Limma as the data was microarray data, using the same Limma pipeline as mentioned above in our FE analysis. As the results were weaker in Murton et al, we chose to use P <0.05 as a cutoff for the Fisher exact test. For the set from Popov et al(dataset 3)(Popov, Makhnovskii et al. 2019), we grouped all the 4h post exercise results vs all 4h control non-exercised and performed DE analysis using edgeR with no other covariates using the same settings as our FE analysis above, with significance based on the found FDR values.

## Results

### Differential Expression analysis

In figure 1 we can see the visualization of the PCA analysis of the count data. There is one main point: there is no clear pattern separating the breeds based on the first two components. Based on the lack of separation of the breeds we gain confidence in the application of a common breed analysis. Any of the lower variance components have a lower proportion of the variation explained than the two observed Principal Components, therefore we are confident that no major proportion of the variation is directly driven by breed. We do observe a significant and detectable effect of breed expression level (as seen further down), meaning there are features in our data which *can* separate the breeds.

**Figure 1.**
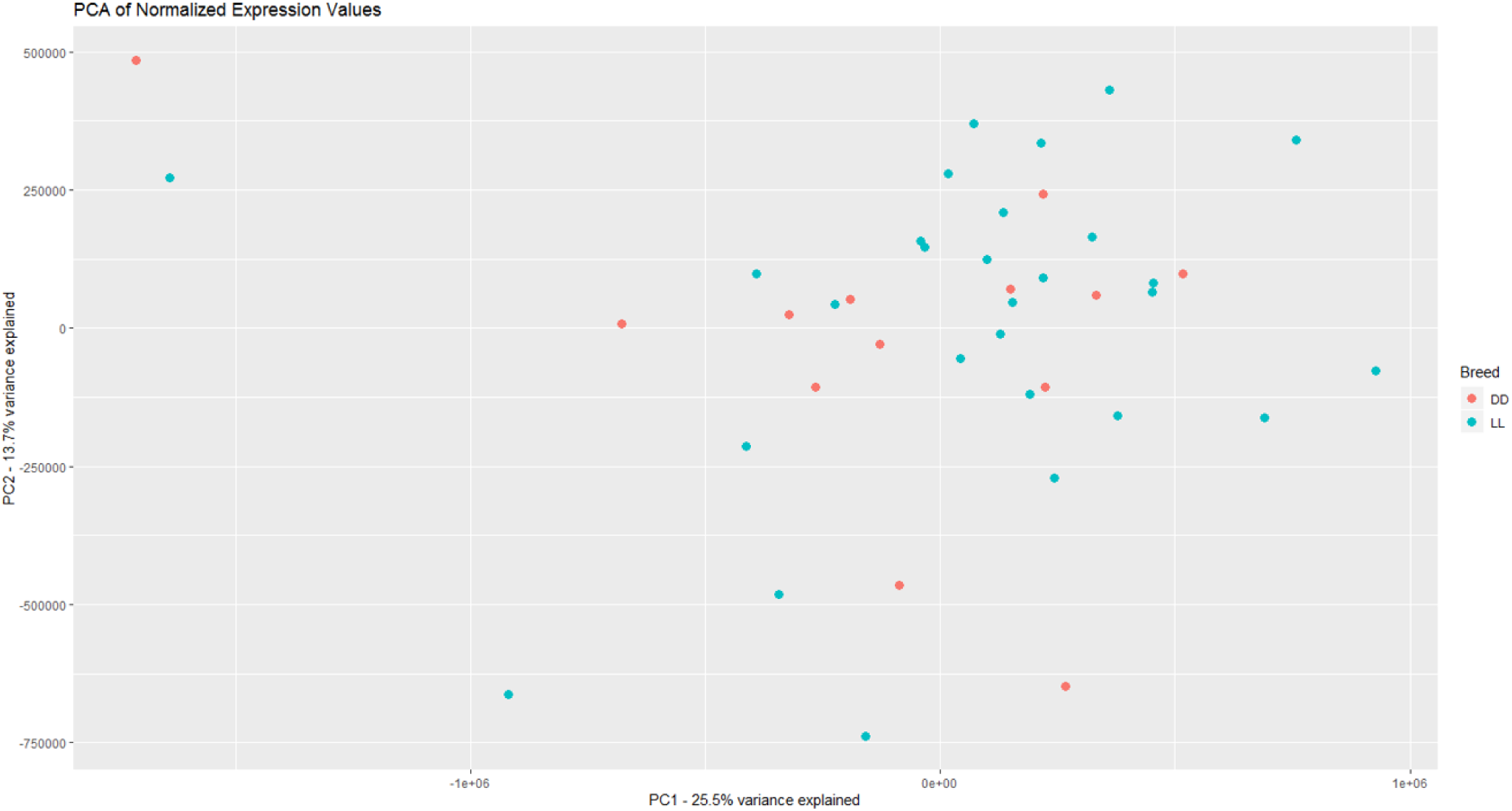
Visualization of the two first principle components in the expression data, with DD being Duroc and LL being Landrace.There is not a clear separation between breeds based on the overall expression, giving credence to a joint breed analysis of the data.

In figure S1 we can see the distribution of the uncorrected p-values for the Deseq2 analysis in our two breeds in relation to FCR with the corresponding figure for the common analysis in figure 2(right). In total, the Landrace analysis had one gene with an FDR < 0.1, and Duroc had 8, and we found 4 in the common breed analysis. Overall, we only find a limited set of genes associated with FCR. In table 1, we see the overview over the genes that where differentially expressed at the 0.1 FDR level in the common and individual breed analysis from Deseq2. As in previous studies, we find genes related to mitochondria (MRPS11, MTRM1) and glucose a related gene (*MGAT4A)(Ohtsubo, Takamatsu et al. 2005)*. We also find genes that have been associated with meat quality phenotypes in cattle and pig (MTRF1,KLH30) (Jiang, Michal et al. 2009, Chung, Lee et al. 2015, Dos Santos Silva, Fonseca et al. 2019). Perhaps the most interesting result, is that one of the genes in the Duroc analysis, TRIM63, has been associated as a biomarker for differences in response to exercise induced muscle damage(Baumert, G-REX Consortium et al. 2018), which ties into our comparison to human data below.

**Table 1.** Overview of genes with a FDR value < 0.1 in all 3 differential expression analysis. There is only a limited amount of genes differentially expressed at 0.1 FDR level for FE. Notably, out of 4 genes in the common breed analysis there are two genes with mitochondrial related Gene Ontologies - MRPS11, MTRM1. MTFR1 has been implicated in eating quality (measures of meat quality post cooking) in cattle(Jiang, Michal et al. 2009) and as a meat PH QTL in pig(Chung, Lee et al. 2015). Also interesting to note that TRIM63 has been suggested as a biomarker for difference in response to exercise-induced muscle damage(Baumert, G-REX Consortium et al. 2018), KLHL30 has been associated with intramuscular fat and muscle metabolism in Nelore Cattle(Dos Santos Silva, Fonseca et al. 2019). MGAT4A has been linked to diabetes and glucose transport (Ohtsubo, Takamatsu et al. 2005).

| Gene Name | Breed | FDR | Regulation |
| --- | --- | --- | --- |
| **PNCK** | Landrace | 0.0007 | Down |
| **Patr-A** | Landrace | 0.08 | Down |
| **MTMR11** | Duroc | 0.07 | Up |
| **C3** | Duroc | 0.02 | Down |
| **LCP1** | Duroc | 0.02 | Up |
| **TRIM63** | Duroc | 0.08 | Down |
| **KLHL30** | Duroc | 0.07 | Down |
| **NANOS1** | Duroc | 0.08 | Up |
| **IGHM** | Duroc | 0.07 | Up |
| **ETV5** | Duroc | 0.02 | Down |
| **MTFR1** | Both | 0.068 | Down |
| **MGAT4A** | Both | 0.098 | Down |
| **SLC38A2** | Both | 0.098 | Up |
| **MRPS11** | Both | 0.067 | Up |

**Figure 2.**
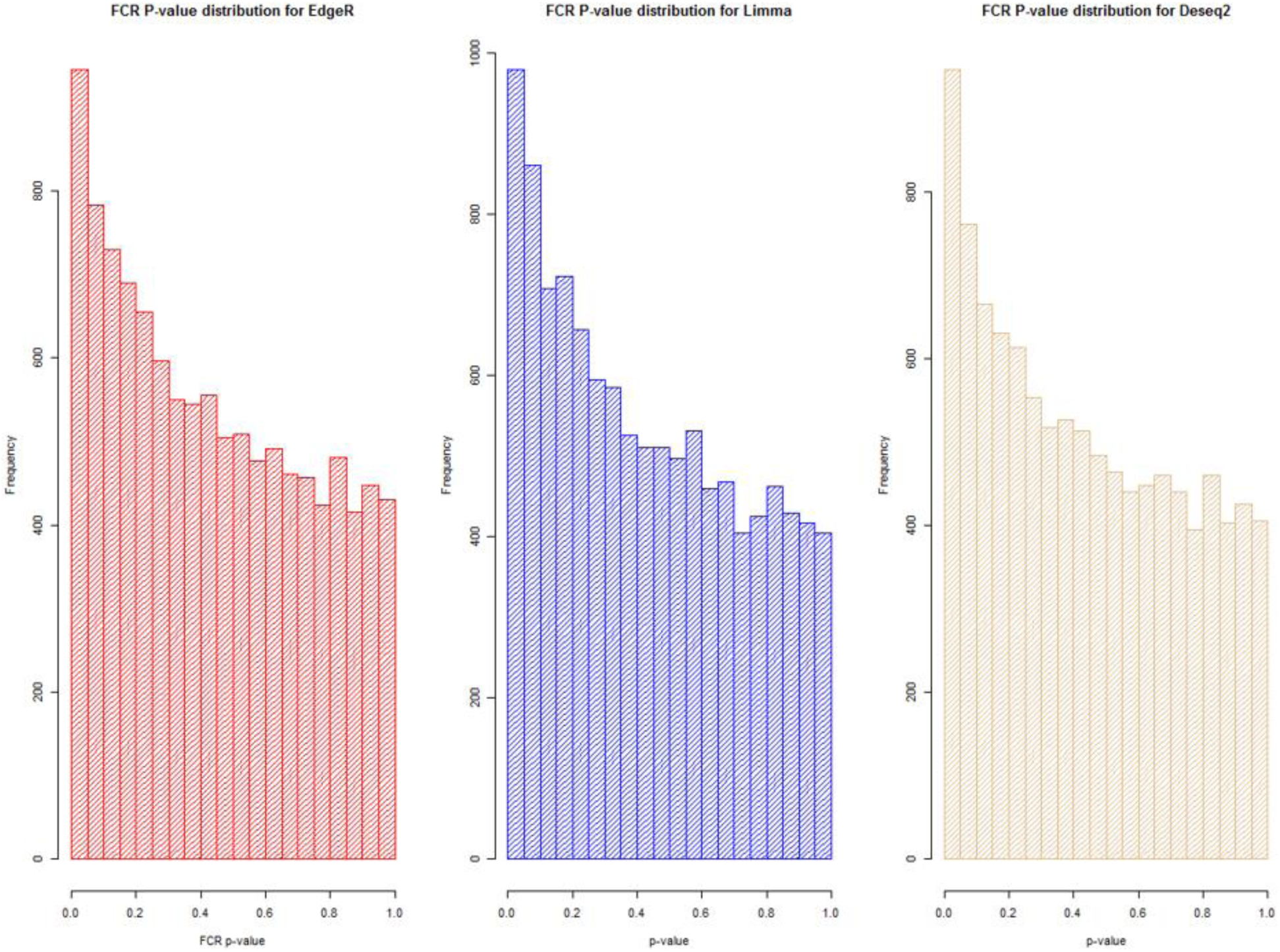
Visualization of the distribution of the p-values testing the relation between FCR and gene expression for all three analysis methods. It is clear in all cases that we observe an anti-conservative distribution, that is, there is an overweight of low p-values.

As the results were somewhat limited, we chose to continue with a different strategy in the joint breed analysis. Based on the results in figure 2, we see that p-values had an overall anti-conservative distribution for FE in the joint analysis, which showed us some promise for further analysis. We chose to calculate the DE using 3 methods, as we wanted to ensure that our results where robust and replicable, knowing that individual methods can vary in output (Seyednasrollah, Laiho et al. 2015). In figure 2 we can see the overview of the distribution of uncorrected p-values for FCR in all 3 methods, showing an anti-conservative distribution regardless of the method. If FCR was unrelated to gene expression in general, we would expect a uniform p-value distribution in our model. We can statistically confirm the likelihood of our observed p-values under the null hypothesis of no relation between expression and FCR using a Kolmogorov-Smirnov test, and in all 3 methods we reject the null hypothesis with (p-value < 10^-16). This leads us to conclude that there is a relation between the muscle tissue expression and FCR. In table 2 we can see the overview over the significance of our covariates in the 3 methods used for DE analysis. The most significant covariate is RIN, highlighting the importance of correcting for the RIN values when analyzing samples acquired in a non-laboratory setting. It has been previously shown that while RIN values do have an impact on expression values, explicitly controlling for this in a modelling framework should appropriately correct the data in most data points(Gallego Romero, Pai et al. 2014). Furthermore, we see that many genes are differentially expressed between the breeds, which is expected, and that age has an impact on expression. To quantify the observed link between expression and FE, we continue with two strategies – analyzing the overall pathway enrichments for the most significant genes and creating gene expression modules based on network analysis of our gene expression profiles.

**Table 2.** Over view over the number of genes with FDR < 0.1 in the common breed analysis for all 3 methods and each covariate. In general, we have modest amount of DE genes for FE, while our other covariates have a amny significant genes associated with them.

| *Trait* | *EdgeR* | *Limma* | *Deseq2* |
| --- | --- | --- | --- |
| ***FCR*** | *4* | *0* | *0* |
| ***Breed*** | *3633* | *3679* | *3428* |
| ***RIN*** | *5572* | *5763* | *5779* |
| ***Age*** | *503* | *189* | *328* |

### Enrichments Analysis

The first step in an enrichment analysis is to select a suitable set of genes. The most general strategy is to pick genes that are differentially expressed after multiple testing correction for such a set. In our analysis, we do not have enough of these for a meaningful enrichment analysis, but we are able to demonstrate an overall relation between FCR and gene expression as seen above in figure 2. In our case, we could select genes with an uncorrected p-value below 0.05, but this is somewhat arbitrary distinction(Butler and Jones 2018). Instead, we chose to make an estimation of the number of additional low p-values in comparison to the uniformly distributed p-values, which represents the null hypothesis of no overall relation between FCR and gene expression. We call this value the divergent count. In essence, we are estimating the interval with the maximum positive divergence between our observed p-value frequencies and the same number of uniformly distributed p-values, assuming an approximately monotonely decreasing p-value distribution in our results. This has the advantage of not relying on arbitrary cutoffs but instead being a property of the overall p-value distribution. In figure 3, we can see a schematic representation of the divergent count. In Figure 4 we can see the a Venn diagram showing the overall divergent counts and overlaps for all three methods, with the full overlap set being the final gene set for enrichment analysis. We can see that a majority of the selected genes are identified by all three methods. This gives us confidence in the robustness of the selected set. To identify enriched functional pathways in our dataset, we chose to use is GOrilla(Eden, Navon et al. 2009). In GOrilla it is possible to give a background set to base the analysis on, making it advantageous for expression data, as it allows us only to use genes actually expressed in our data as a background. For the full output of the analysis, see supplementary table 2. Overall, 5 terms were significant post multiple corrections, with 4 out of these being related to mitochondrial ontologies In figure 5 we can see a summarized output of the significant post multiple testing correction GO-terms and groups based on the GOrilla analysis, using Revigo(Supek, Bosnjak et al. 2011). Based on this, the important overall pathway was translation elongation.

**Figure 3.**
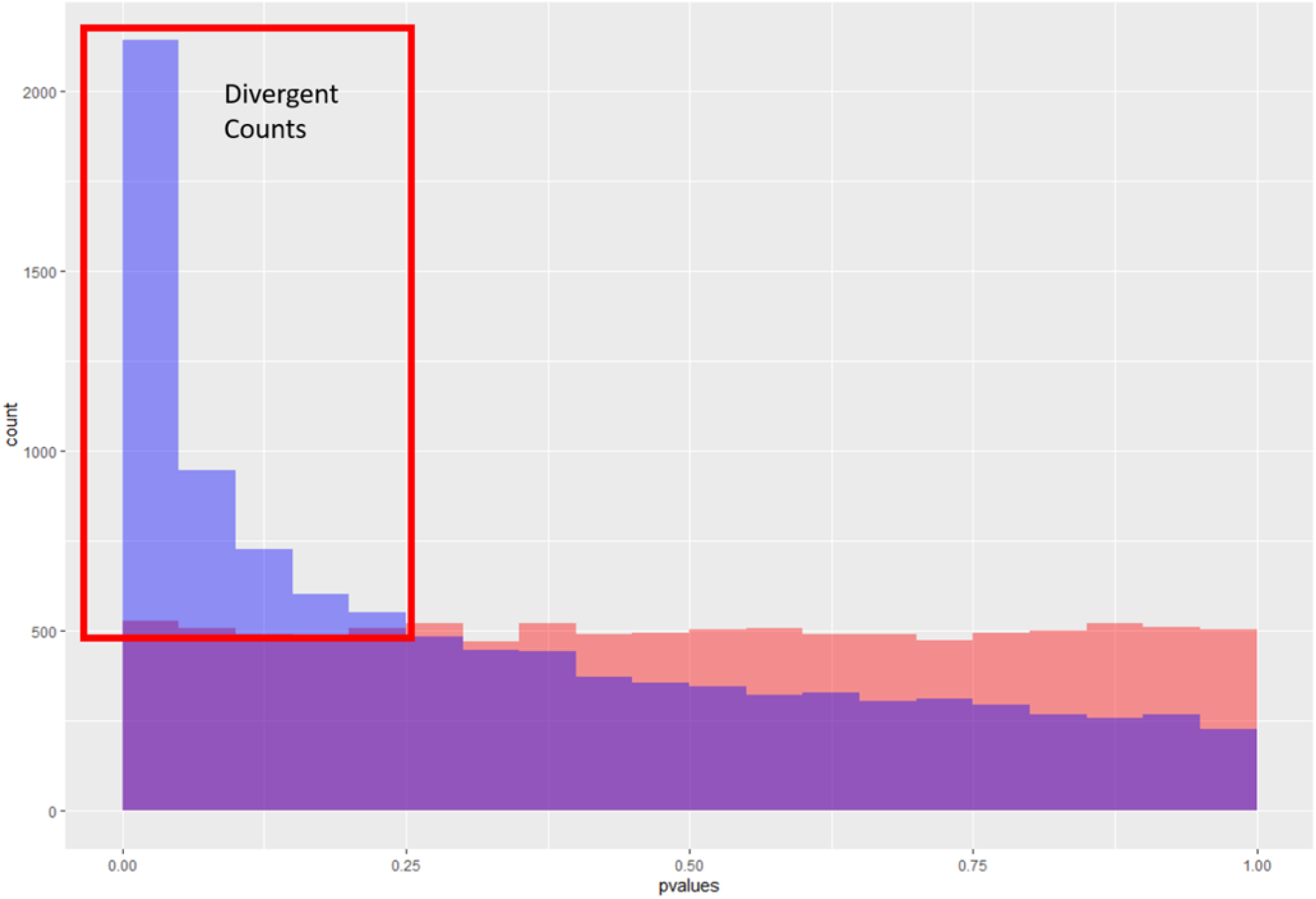
Schematic representation of the divergent counts. Here we see to theoretical p-value distributions, one which is uniform (in red) and one which is anti-conservative (blue). The purple area is where they overlap, and the blue area is the area used to estimate the divergent counts.

**Figure 4.**
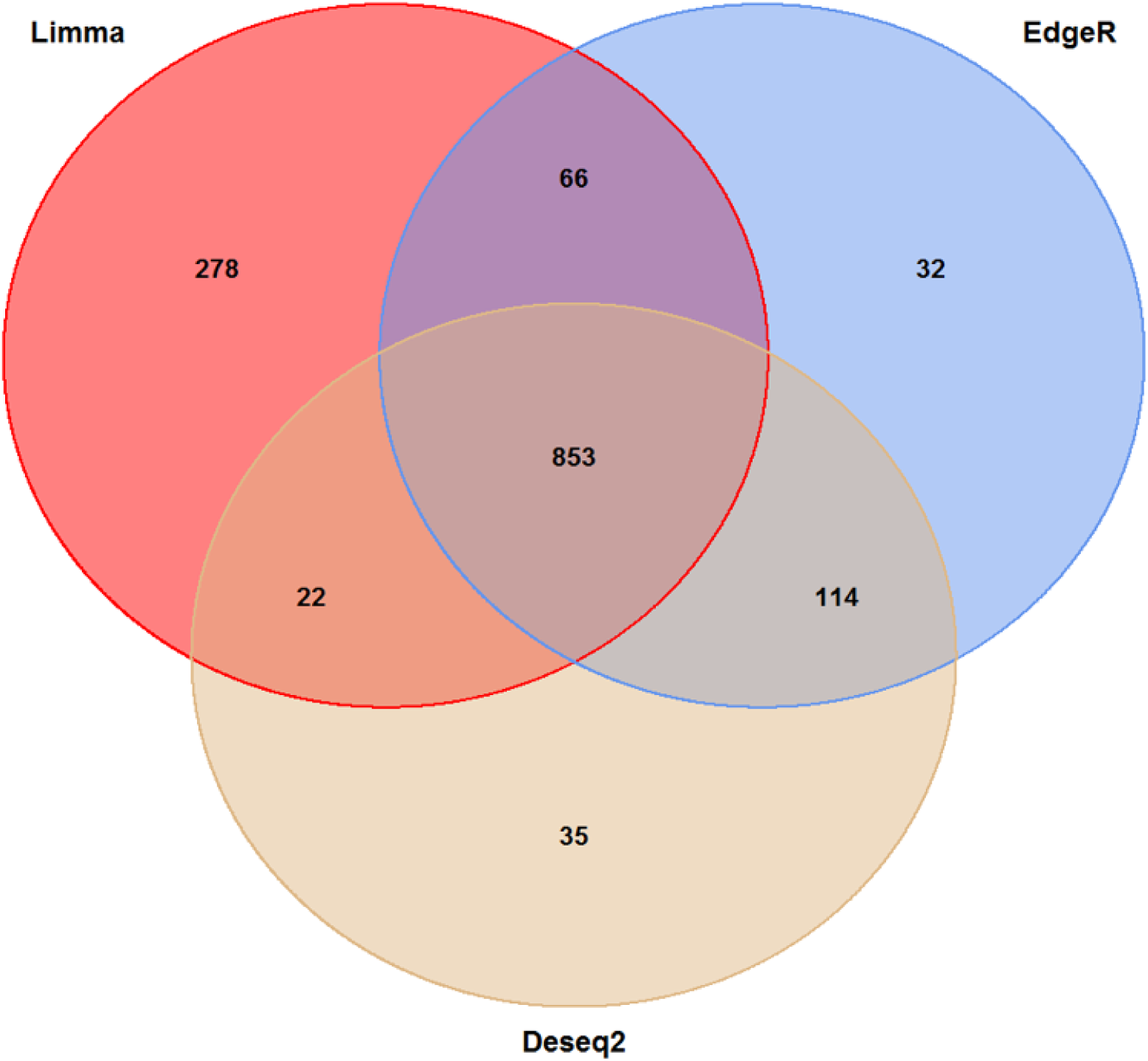
Venn diagram of the overlap in the divergent counts between the three methods. We see here that the Limma is overall less conservative than the two other methods, but in general, the methods are in high agreement with each other. The final set of genes selected for the enrichment analysis was the 853 triple overlapping set.

**Figure 5.**
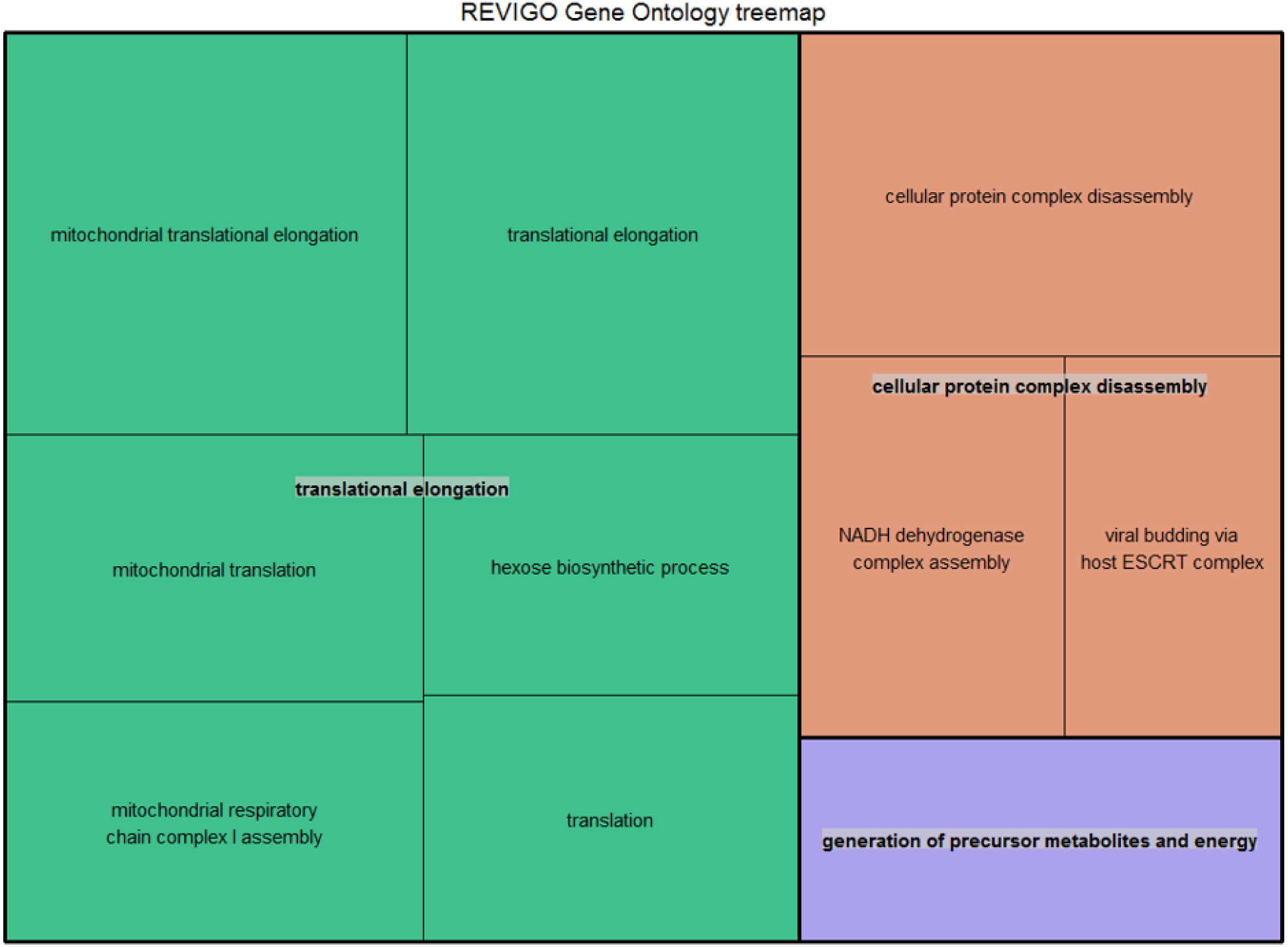
Summarized representation of significant GO- for the genes set generated from the divergent count (853 total genes) overlap based from the DE analysis of FCR. The size of the boxes is scaled according to the −log10 of the p-value. The most significant individual terms are all in the translation, indicating a link between mitochondrial activity and FE.

### Gene Interaction Analysis

Many strategies can be used to take advantage of the interaction or co-expression between genes. We propose to apply modelling of pairwise gene interactions, which explicitly includes the phenotype of choice, which in our case is FE. This can be advantageous when dealing with complex phenotypes, as it may allow us to capture subtle biological variation. We chose to perform the gene interaction analysis based on the set of genes we identified from the divergent counts in our DE analysis. The visualization of the empirical divergent counts and the bootstrapped counts can be found in supplementary figure 2. Based on these results, the maximum bootstrapped divergent count was 83, and we observed 193 genes with a divergent count over 83. This means that many of the genes’ p-value distributions are very anticonservative, and not very likely to happen by chance. There is however, the issue of data independence, as the genes’ results are not independent from each other. Due to this, and general concern of data size and weak effects we used a conservative qualitative heuristic and focused on the top 20 genes based on our methodology. From the top 20 genes (see supplementary data 3 for the full results), the overall highlights were several transcription regulators: ETV1(an androgen receptor activate gene), LF1 (transcription factor) and KDM4C (transcription activator and growth related gene) (Bray and Kafatos 1991, Cai, Hsieh et al. 2007, Gregory and Cheung 2014); two mitochondrial genes, KMO and MRPS11(Meinke, Kerr et al. 2019),; two genes related to muscular atrophy - GEMIN7 and PLPP7 (Baccon, Pellizzoni et al. 2002, Meinke, Kerr et al. 2019); on gene implicated in heart development BIN1 (Nicot, Toussaint et al. 2007), two lipid metabolism/obesity related genes ACOT11 and GPD1 (ADAMS, CHUI et al. 2001) (Park, Berggren et al. 2006); and finally 3 genes associated with specific traits in pig IL2RG (Immune system in pigs)(Suzuki, Iwamoto et al. 2012), GGPS1 (meat quality) and PPARA (weak association with fat percentage) (Szczerbal, Lin et al. 2007). Interestingly, MRPS11 was also differentially expressed.

### Gene Network Analysis

Based on our network analysis, we identified 19 distinct modules after correcting for RIN and merging the modules based on similarity. Based on the DE analysis, we decided not to focus individually on Landrace or Duroc pigs in the network analysis, and thus the network was generated combining both breeds. Looking at the the clustering in figure 6a, initially one might think that the network is poorly constructed, as the module dendrogram representation is not very clear. In general, we see that some modules look closely clustered based on the dendrogram, such as the red module, while other are more diffuse. We should however realize that the modules themselves are based on N x N matrix, where n is >10.000. Thus, it is not easy to represent the modules properly in lower dimensions. Therefore, we rely on the module eigenvalue trait correlation and pathway analysis of the modules to asses if they are biologically meaningful. In figure 6b we can see the correlation between the eigenvalue of the modules and the traits and covariates we included in the DE analysis. We observe that the RIN correction of the individual genes has removed all the effect of the RIN on the eigenvalues of our modules. Several of the modules are well correlated with the breed and age, with correlation > 0.5, while FCR is mainly correlated with two modules, red and turquoise. The red and turquoise module include 391 and 3744 genes, respectively. Based on these results we performed GO-term analysis on the red and turquoise module. The red module is more correlated to breed and age than FCR, but we know that breed and FCR are correlated, and in our data, age is correlated with FCR (0.5). It should be noted that the age and FCR correlation is caused by the higher FCR pigs in our data exhibiting lower growth rates, thus needing more time to reach the tissue sampling as the slaughter takes place at a target weight of approximately 100 kg. The turquoise module shows highest correlations in FCR. In figure 7 we see the Revigo summary of the GOrilla GO term analysis performed based on the genes in the red (a) and turquoise (b) modules. In both the red and turquoise modules, a large number of GO terms where significantly overrepresented after multiple testing correction (see supplementary data 4 and 5 for the full list for red and turquoise respectively), indicating that the modules do represent specific biological pathways. In the red module, the most significant group of terms where related to mitochondria, which were grouped into three overall groups – translation elongation, electron transport chain and hydrogen ion transmembrane transport. This mirrors our finding from the DE analysis and the gene interaction analysis. As the module has a negative correlation with FCR, it indicates a relation between higher mitochondrial activity and lower FCR, thus higher efficiency. In the turquoise module, there was one large grouping of terms – DNA repair. This category included many GO terms, related to RNA, DNA, animo acid and nucleic acid metabolism and processing. These processes could be seen as generic growth and maintenance processes, and as the module is positively correlated with FCR, we can speculate the higher activity in DNA repair and related processes are increasing energy spend on maintenance, thus lowering efficiency. Due to the size of the module and the processes involved, it seems that the turquoise module is generically associated with overall cell maintenance and growth processes, giving it a somewhat unspecific functionality. In supplementary data 6 we find the top 10 most connected genes in the red and turquoise module. Interestingly, in the red module 7 out of 10 genes belong to the NADH ubiquinone oxidoreductase group (NDUF), with the remaining 3 also being implicated in mitochondrial function. Thus, the mitochondrial genes are both overrepresented in the red module and the most connected within the module. In the turquoise module, the results are unclear, as the most connected genes do not belong to any specific process, but instead cover a range of general processes that are generally important for cell function. This agrees with the general observation based on the size of the module and the overrepresented GO terms.

**Figure 6.**
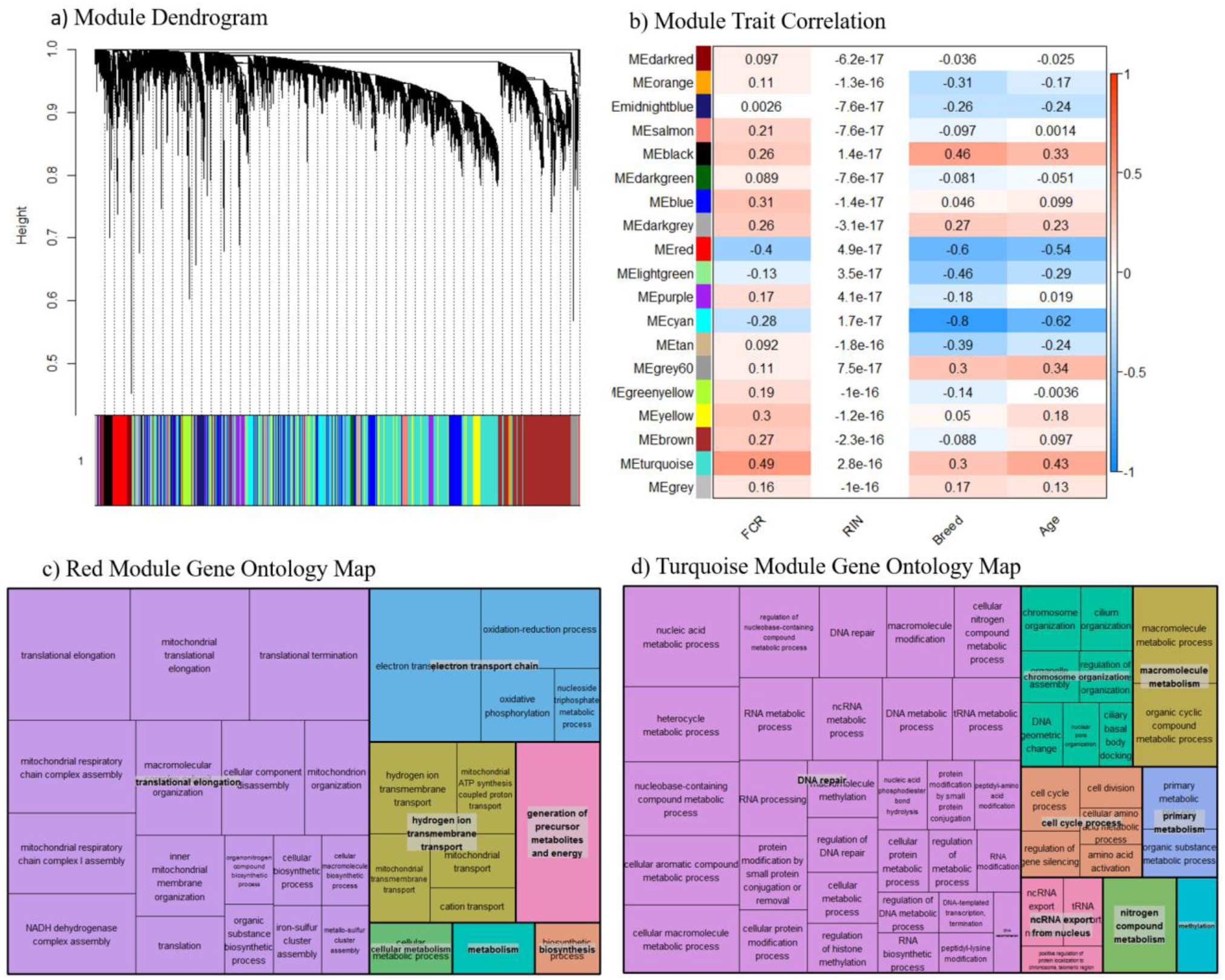
(a) Dendrogram over the module clustering. Looking at the visual clustering not all the modules look equally well defined, but it should be noted that the actual relations in given module cannot be simplified to two dimensions, as the all the relations between the genes exist in N dimentional space, where N is the number of genes. (b) Correlation between module eigenvalue and our traits, including RIN. We see here that the correlation to RIN is essentially 0 in all cases, indicating our linear correction method has worked well. Based on the top two modules **(c)** Summarized representation of significant GO- for genes in the red module of the WGCNA network analysis. The three largets groups are all associated with mitochondria, mirroring the results found in the differential expression analysis and the gene interaction analysis. (b) Summarized representation of significant GO- for genes in the turquoise module of the WGCNA network analysis. The main grouping here is DNA repair, which is not found in our other analysis. This may represent that increased energy expenditure on maintenance processes is reducing FE.

**Figure 7.**
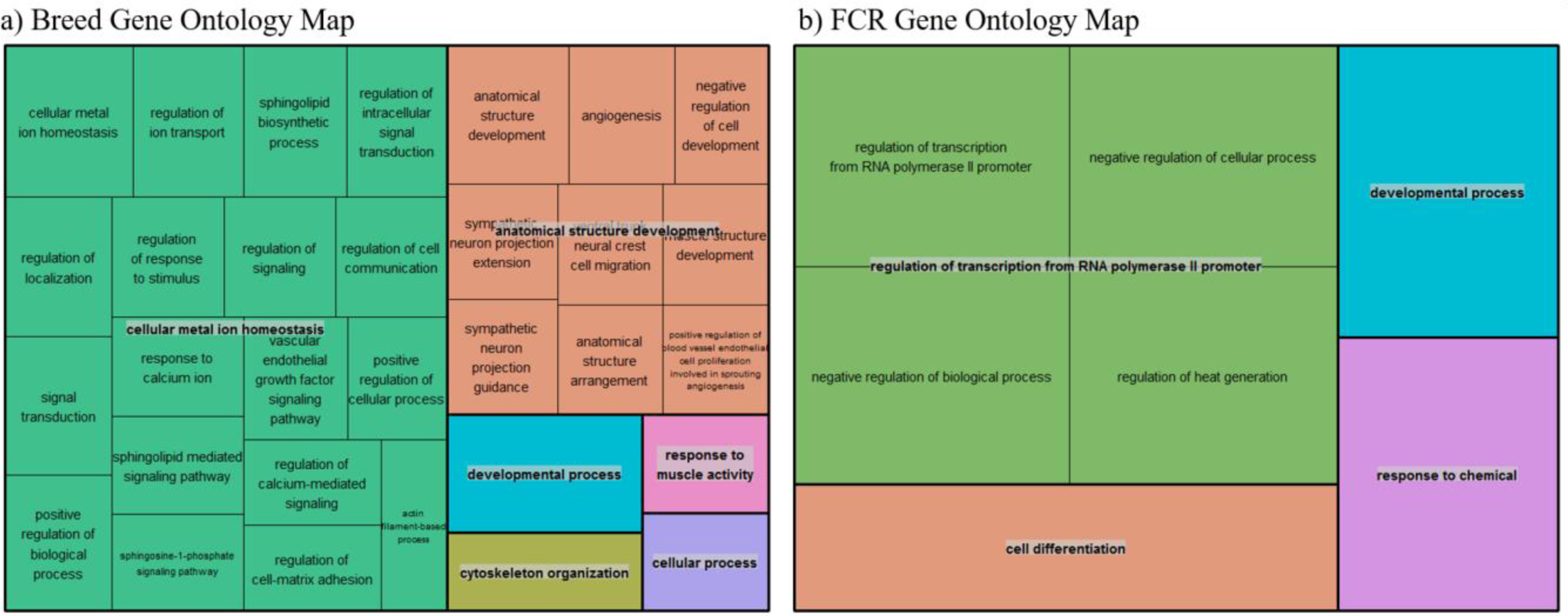
(a) Summarized representation of significant GO- for genes significantly associated with exercise in one of the three human dataset and between the breeds, based on a total of 702 genes. The size of the boxes is scaled according to the −log10 of the p-value. Here we find two overall main categories, cellular metal ion homeostasis and anatomical structure development. (b) Summarized representation of significant GO- for genes significantly associated with exercise in one of the three human dataset and in our divergent set for FCR. The size of the boxes is scaled according to the −log10 of the p-value. Here the main process is regulation of transcription from RNA polymerase. Overall, the categories are not very significant here as it is only based on 42 genes.

### Human Exercise Data

To test they hypothesis that improvements in efficiency could be linked to a state mimicking exercise, we compared our divergent counts genes for FCR and the genes differentially expressed between breeds with 3 different human exercise datasets [33-35]. The results can be found in table 3. We are comparing if there is a higher proportion of genes that are significant for exercise-mediated changes in our two subsets, breed and FCR related genes, in relation to the non-differentially expressed genes. We see that in all cases there is a higher proportion of significant genes in the breed and FCR set versus the background set, as the odds ratio between the subsets and the background is always below 1. In general, the breed results are more significant than the FCR genes, but they show similar ratios. This is likely because there are roughly 4 times more breed genes, yielding higher statistical power. Given the overall results, it does seem like both FCR and breed related genes are slightly more significant than background for exercise related changes. We also did the pathway enrichment analysis for the genes that where significant in both one of our three human data sets and in the breed, and FCR set respectively. The overall results are found in figure 7a (breed) and 7b(FCR). In the breed, we find that main categories are cellular metal ion homeostasis and anatomical structure development, based on 702 genes. For FCR, only 42 genes overlap with the human significant genes, meaning the results of the enrichment are not as significant, but the main overall group is regulation of transcription from RNA polymerase II promoter.

**Table 3.** Results of Fisher exact test comparing the number of genes significant for difference in rested and exercised muscle in divergent count genes for genes found in the divergent count for FCR and breed and the background for each of the 3 human data sets(dataset 1 (Devarshi, Jones et al. 2018),dataset 2 (Murton, Billeter et al. 2014) and dataset 3 (Popov, Makhnovskii et al. 2019)).

| **Data** | **P-value Breed** | **Odds ratio Breed** | **P-value FCR** | **Odds ratio FCR** |
| --- | --- | --- | --- | --- |
| **Dataset 1** | 0.0017 | 0.79 | 0,0046 | 0.71 |
| **Dataset 2** | 0.0012 | 0.85 | 0.22 | 0.9 |
| **Dataset 3** | 0.12 | 0.84 | 0.47 | 0.88 |

## Discussion

There have been 4 previous studies analyzing the muscle transcriptome in an FE context (Jing, Hou et al. 2015, Vincent, Louveau et al. 2015, Gondret, Vincent et al. 2017, Horodyska, Wimmers et al. 2018). The study by Gondret et al [18] was based on selecting divergent FE lines of Large White pigs for 8 generations, used 24 samples and was based on microarray. They reported a high number of differentially expressed genes in muscle between the low and high RFI groups (2417), but it is not clear from their paper how many probes were included in the statistical analysis and how this may affect multiple testing correction. They also reported that a gene was considered differentially expressed if one probe met the cutoff regardless of multiple probes did not. They reported that mitochondrial electron chain transport, glucose metabolic process and generation of precursor metabolites and energy as significant pathways for RFI.

In the study from Horodyska et al [17], they used 16 pigs, but included 8 pigs of each gender. They used an uncorrected p-value of 0.01 as their threshold,, with no consideration weather this is appropriate given their overall data distribution. They report 272 genes with p-value < 0.01, which is similar to ours of 243, however we have included less genes in our analysis (14497 vs 10563). Overall, we cannot assess their results as very significant.

In Vincent et al [20], they had 16 female Large Whites from divergent RFI lines, their study was microarray based, but they reported their results based on uncorrected p-values in both expression and proteomics. They do however report finding mitochondrial related probes being significant.

Finally, in Jing et al [19], they had a total sample size of only 6 Yorkshire pigs, based on the selection of the most extreme RFI pigs in a set of 236. They reported 645 DE genes, with 99 with FDR lower than 0.05. However, selecting such few samples at the extreme end of FE does raise the question of replication, as the large differences in RFI/FCR they achieved could easily be caused by factors that are not generally applicable. They found that the most significant pathways in their data were mitochondrial activity, glycolysis and myogenesis pathways. Despite the issues presented with the studies, it is notable that mitochondria are reported to be related to FE multiple times.

In our study, we have the highest number of samples reported (41) and we include two breeds, which do not have directly divergent selection for FCR, but with one of the breeds more positively selected for FCR. Having this setup does present advantages and disadvantages. The advantage in relation to the other studies it that the results may generalize better across breeds. The disadvantage is that we may be fitting breed effects instead of phenotypic effects, but we do account for breed in all our analysis. The other main difference is that we have selected pigs with a range of FCR values, and fit FCR as a continuous value. In general fitting a continuous value is more informative, and the fact that we have a range of pigs that are not divergently selected, may make the results more applicable to a real life setting. In pig production there is no low FE selected line to contrast with, so the biological background of FE in a normal breeding population may be more relevant and interesting.

Another general issue is how to deal with statistical issues in analysis of FE. From the various studies presented above it is clear that FE is a somewhat subtle phenotype in muscle tissue, and thus a lot of data is needed make conclusions. Here we try to tackle this issue by not being overly conservative, but still applying multiple testing correction by using and FDR of 0.1 level for individual results in our DE analysis. Furthermore, we generally try to analyze our data by either taking the overall distribution of results and/or combing genes in groups, to avoid relying on individual weak results.

### Differential Expression analysis and Pathway Enrichment

We have analyzed the transcriptomic differences and molecular pathways involved in differences in FCR in two different breeds. Based on DE, we identified 14 genes with an FDR value below 0.1. The highlights here were the finding of mitochondrial genes, and TRIM64, which related to exercise induced muscle damage.

Due to the limited results in the DE analysis, we chose to use a novel approach to perform a pathway enrichment analysis. In practice, we wanted to broaden the number of genes for the pathway analysis, but at the same time also select a robust and meaningful set of genes. To make the analysis more robust, we choose to base the pathway analysis on results from 3 DE expression methods. Furthermore, we elected to select genes based on the overall divergence from the null hypothesis of our p-value distribution, as this should represent a set of genes that is likely to be associated with our trait, even the genes are not significant based on individual FDR corrected p-values. To our knowledge, this is a novel way of selecting a group of genes, which we called the divergent count. Looking at the enriched pathways in our dataset selected based on the divergent counts, we find results that are common in the literature in several species beyond the pig studies already mentioned(Connor, Kahl et al. 2010, Bottje, Lassiter et al. 2017), namely differences in mitochondrial pathways related to FE, summarized as mitochondrial translation elongation in our Revigo summary. While this is not a novel result, we did find it in a novel setting, with larger sample size, novel population selection and using a continuous value for FCR. This acts as further evidence to the link of mitochondrial activity and FE, but also as evidence that it may be relevant in real breeding populations, and not only in divergently selected test populations.

### Gene Expression Interaction

Our gene expression interaction analysis is a novel way of finding the most important genes, which has not been applied to FE in pigs before. Based on the qualitative analysis of the top 20 genes, the results seem promising. We found several transcription factors, including the most divergent gene (ELF1), which makes sense in regards to gene interaction. The remaining genes also seemed promising, as they included categories one can expect to be related to muscle growth and FCR, such as lipid metabolism and muscle atrophy. Confirming previous results, we also identified two mitochondrial genes among the top 20.

### Gene Network Analysis

Our gene network analysis revealed two modules with a correlation > 0.4 with FCR. Based on the GO term analysis enrichment of the red module, we find many enriched GO terms related to mitochondrial processes, confirming our finding in the other analysis, and from other studies. More specifically, the negative correlation between the red module eigenvalue and FCR also shows that higher mitochondrial activity is positively associated with higher efficiency. Based on the top ten hub genes in the red module we confirm this picture, as all ten genes are related to mitochondria, and seven of them are from the NDUF family, which was also found in the gene expression interaction analysis. The turquoise module was the most correlated module(0.49), and furthermore, it was more correlated to FCR than to our other traits. Based on the GO term analysis, we found that the cluster was highly enriched for genes related to DNA repair, which included GO terms relate to RNA, DNA, animo acid and nucleic acid metabolism and processing. To the best of our knowledge, this is the first evidence of these processes being related to FE in general. The only previous link to DNA repair in livestock was a feed restriction study of cattle(Connor, Kahl et al. 2010). The top ten hub genes of this module did not show a clear picture, with the genes involved in a wide range of processes related to general cell maintenance. This indicates that the turquoise module represents general housekeeping functions, rather than very specific pathways. As the module eigenvalue was positively correlated with FCR, we can speculate that more active DNA repair and maintenance processes represent higher maintenance costs, thus reducing efficiency.

### Human Exercise

We have established earlier that the gene expression and molecular background of FE is still somewhat elusive. To try and identify what overall mechanisms could be at play, we hypothesized that differences between our two breeds, which have different overall FE, and genes related to FCR, are more likely to be important for processes involved in exercise. The reason we had this hypothesis, is that the pigs are selected for lean growth, and it is possible that this growth stimulus is similar to the effects induced in muscle by exercise. We found a slight confirmation of this hypothesis, as we found similar favorable odds ratio for our hypothesis in all 3 datasets we tested for both FCR and our breed genes. Our pathway enrichment analysis for FCR did not yield any very significant results, as it was only based on 42 genes. The main overall category identified, based on 4 go terms, was regulation of transcription from RNA polymerase II (pol II) promoters. Interestingly, Actin has been associated with the pre-initiation complex necessary for transcription by RNA polymerase II(Hofmann, Stojiljkovic et al. 2004), which could be relevant given the importance of actin in muscle tissue(Tang 2015). There are also links between a poll II subunit and myogenesis (CORBI, PADOVA et al. 2002). Although these results may be relevant, our data here is too weak for solid conclusions.

In regards to the genes overlapping between exercise and breed differences, the results are more statistically robust, as they are based on an overall larger gene set of 702 genes. Here we find two overall groups – cellular metal ion homeostasis and anatomical structure development. For the first term, we know that the transport of ions is generically vital to muscle function (Wolitzky and Fambrough 1986, Mohr, Krustrup et al. 2007). The second overall term, anatomical structure development, is very generic in terms of function, and includes sub-categories that are related to muscle development, such as muscle structure development.

Overall, the results from the Human data analysis represent a novel hypothesis, but requires more analysis and new experiments on pigs to strengthen the link between FE and exercise. One interesting aspect of this analysis is that in theory pigs could be used as a model for lean growth in sedentary conditions, which in the long run could yield interesting therapeutic possibilities applicable to humans.

## Conclusion

We have analyzed the muscle transcriptome from Duroc and Landrace, twp of the main purebred breeding pigs in Denmark. In contrast to previous studies, we did not use any lines divergently selected for FE, and we included a wider range of FE values, which were modelled as a continuous trait, using the highest number of pigs in a study of this type. We identified several individual genes based on DE analysis and gene-gene interaction analysis that are involved in FCR, with many of them having relevant functional backgrounds from previous studies. We applied a novel strategy to select genes for pathway enrichment, the divergent count. Based on enrichment analysis, gene-gene interaction, network analysis and DE we found several interesting candidate biomarkers genes and pathways. We reinforced the knowledge that mitochondrial activity is important FCR, but using a non-divergently FE selected pig population. Based on the findings, we postulate that mitochondrial genes, and in particular genes from NDUF group or MRPS11 could be used as potential biomarkers for FCR in pigs. Furthermore, all our top genes from our interaction analysis also show promise as potential FCR biomarkers. Finally, we find that there is a putative link between genes involved in exercise related changes in human, and FE in pigs

## Ethics

As animals were only sampled post conventional slaughter, no ethics approval was needed for the study.

## Author Contributions

HNK conceived and designed the project and obtained funding as the main applicant. VAOC and HNK designed the muscle sampling experiments, phenotype data collection and statistical/bioinformatics analyses. VOAC performed the sampling, data processing, data visualization and bioinformatics and statistical analysis. All authors collaborated in the interpretation of results, discussion and write up of the manuscript. All authors have read, reviewed and approved the final manuscript.

## Acknowledgments

We would like to thank all group members of the Quantitative Genomics, Bioinformatics and Computational Biology Group for participating in discussion when needed and creating a fruitful scientific environment.

## 1 Data Availability Statement

The data will be uploaded to GEO and released if the article is accepted

